# Infect While the Iron is Scarce: Nutrient Explicit Phage-Bacteria Games

**DOI:** 10.1101/674382

**Authors:** Daniel Muratore, Joshua S. Weitz

**Affiliations:** School of Biological Sciences, Georgia Institute of Technology, 950 Atlantic Drive, Atlanta, GA, USA 30332; School of Physics, Georgia Institute of Technology, 837 State Street, Atlanta, GA, USA, 30332

**Keywords:** Game Theory, Viral Ecology, Siderophores, Marine Ecology

## Abstract

Marine microbial primary production is influenced by the availability and uptake of essential nutrients, including iron. Although marine microbes have evolved mechanisms to scavenge sub-nanomolar concentrations of iron, recent observations suggest that viruses may co-opt these very same mechanisms to facilitate infection. The “Ferrojan Horse Hypothesis” proposes that viruses incorporate iron atoms into their tail fiber proteins to adsorb to target host receptors. Here, we propose an evolutionary game theoretic approach to consider the joint strategies of hosts and viruses in environments with limited nutrients (like iron). We analyze the bimatrix game and find that evolutionarily stable strategies depend on the stability and quality of nutrient conditions. For example, in highly stable iron conditions, virus pressure does not change host uptake strategies. However, when iron levels are dynamic, virus pressure can lead to fluctuations in the extent to which hosts invest in metabolic machinery that increases both iron uptake and susceptibility to viral infection. Altogether, this evolutionary game model provides further evidence that viral infection and nutrient dynamics jointly shape the fate of microbial populations.

## 1 Introduction

Photosynthetic marine plankton account for approximately half of global annual carbon fixation [14]. Trace metals, including iron, are necessary for the molecular machinery which perform such important metabolic processes as photosynthesis and nitrogen fixation [28, 39]. However, concentrations of iron in the open ocean surface are sub-nanomolar [3]. Furthermore, due to iron’s low solubility in water and the relatively high concentration of weakly iron-binding organic molecules dissolved in seawater, iron in seawater is largely confined to the particulate phase, either as ferrihydrite minerals or bound to organic particles [28, 37, 50]. Neither of these forms of iron are readily available for uptake by bacteria, rendering iron a scarce resource for photosynthetic picoplankton [20, 22]. Iron limitation has been implicated as the cause of high nutrient low chlorophyll zones (HNLCs), including the equatorial Pacific and Southern Oceans [32, 5]. Iron amendment experiments (such as those reported in [10, 46, 26]) have found that added iron can increase primary productivity in these regions. In contrast, other iron amendment mesoscale experiments have found no release from iron limitation [48], or, interestingly, in the case of the 1995 IronEx II experiment, possible competition for the added iron between diatoms and heterotrophic bacteria [11].

Microbes in host-pathogen, soil, and marine systems have adapted to modulate local iron conditions through the evolution of siderophores [41, 38, 49]. Siderophores are small molecules with much higher iron-binding affinity than ambient dissolved organic matter [8]. Iron, once bound by a siderophore, remains in the dissolved size class. Bacterioplankton can take up the iron-siderophore complex with a corresponding siderophore-uptake receptor. Notably, siderophores can be freely excreted into the environment. Extracellular excretion poses challenges for understanding the benefits of siderophore production for cellular fitness. Apart from wasted energy due to diffusive loss of siderophores, there is an additional “public goods” problem [21]. Cells which express siderophore receptors but do not produce siderophores of their own may have a fitness advantage over producers [7]. The proliferation of “cheating” phenotypes may lead to a tragedy of the commons scenario, by which the siderophore producing population is driven to extinction by competition from cheaters, decreasing the fitness of cheaters by reducing the bioavailable iron pool [47]. Theoretical evolutionary models exploring this problem find that diversification of siderophore structure and corresponding uptake receptors can mitigate pressure from cheating phenotypes [30]. Diverse siderophore structures found in marine systems support the hypothesis that siderophores are a competitive adaptation [3, 8].

Despite their benefits, siderophore surface receptors may also provide a port of entry for viral infection [4]. Some siphoviruses of *E. coli* have been found to use ferrichrome (a siderophore) uptake protein FhuA as a receptor [2]. The T4 Myovirus of *E. coli* uses outer membrane protein OmpC, a different receptor protein which also has metal-binding properties [1]. However, the receptors used by marine viruses remain largely elusive. Comparing the amino acid sequences of tail fiber proteins from marine viral genomes to those of viruses with known receptors is one way to explore the possible catalogue of marine virus receptors. Via a comparative approach, Bonnain et al. [4] found dual histidine residues in marine viral tail fiber sequences homologous to those of *E. coli* phage T4. Structural studies of the T4 tail fiber protein show a small number of iron ions bound to those dual histidine residues [1]. A more recent study of environmental metagenomes and metatranscriptomes from the TARA oceans expedition [9] found extensive evidence of iron-binding residues in phage tail amino acid sequences, with dual histidine residues homologous to T4 and other markers of putative iron binding in 87% of viral tail proteins. These findings underscore the ‘Ferrojan Horse Hypothesis’, according to which abundant viruses of marine picoplankton [15, 44] incorporate iron into their tail fiber proteins to facilitate infection via siderophore uptake receptors. As a result, cells in severely iron-limited conditions, such as cells in HNLCs, may experience a tension between the benefits of nutrient uptake and the danger of viral infection [27].

The Ferrojan Horse hypothesis poses an interesting eco-evolutionary dynamics question: under what circumstances is it evolutionarily advantageous for plankton to produce and take up siderophores? Further, does pressure from viral infection have an impact on the evolutionary dynamics of siderophore production? Previous studies modeling the eco-evolutionary benefits of siderophore utilization [30, 24] have not incorporated feedback due to viral infection. Here, we propose a bimatrix replicator dynamical model of host-virus interactions coupled to an exogenous resource (in this case, iron). In doing so, we assess the ways in which conflicting pressures between resource limitation and viral infection can influence host uptake strategies. As we show, when resource levels are variable, viral infection can drive oscillatory dynamics involving multiple virus types (including some that incorporate iron into their tails) and multiple host types (including some that produce siderophores).

## 2 Results

### 2.1 Evolutionary Game Theory of Ferrojan Horse Hypothesis

#### 2.1.1 Replicator Dynamics for Bimatrix Games

We first construct a non-resource explicit bimatrix game in which hosts compete with viruses (Figure 1). In this game, each individual utilizes one of two phenotypic strategies. Host strategies include producing siderophores (Cooperation) or not (Defection). Virus strategies include iron-tail fiber incorporation (Ferrojan) or not (Non-Ferrojan). The payoff matrix is:

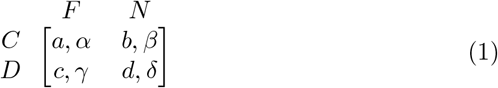

**Fig. 1:**
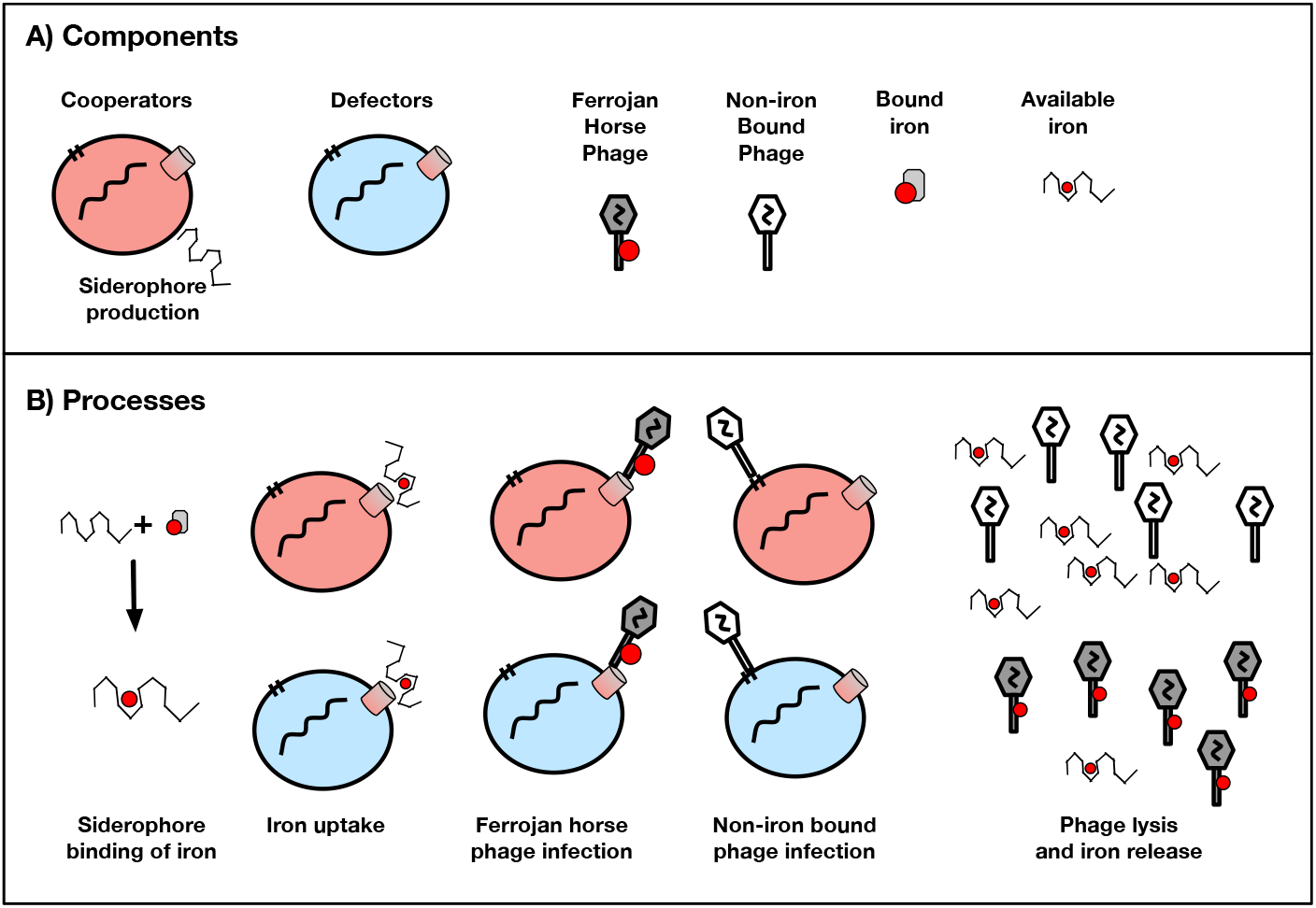
Schematic of bimatrix replicator dynamics in Ferrojan Horse game. (**a**) Phenotypic profiles of hosts and phage. Cooperator hosts produce siderophores, which ligate to biologically inert iron. This allows the iron to be taken up via a specialized receptor (red cylinder). Defector hosts have the receptor, but do not expend energy or resources to produce siderophores. Ferrojan phage have some amount of iron (red circle) bound to their tail fibers, unlike Non-Ferrojan phage. Environmental iron exists in two states - biologically unavailable and biologically available (siderophore-ligated). (**b**) Processes modeled in feedback-coupled replicator dynamics. Siderophores improve local iron bioavailability. Iron ligated by siderophores can then be taken up by either cooperator or defector hosts to grow. However, Ferrojan phage bind to the siderophore receptors to initiate infection. Non-Ferrojan phage bind to some other receptor not involved in iron uptake. When a cell infected by a Non-Ferrojan phage lyses, intracellular iron is released into the bioavailable pool. When a cell infected by a Ferrojan phage lyses, however, some proportion of the iron that would otherwise be in the labile lysate is instead sequestered in phage particles.

We use a bimatrix convention for payoffs: e.g., if a cooperator host (C) encounters a Ferrojan virus (F), the payoff (average fitness) to the host will be *a* while that to virus will be *α*. Table 1 summarizes the relative fitnesses of all strategy combinations in terms of payoff parameters. Using the “battle of the sexes” model as described in [19], the replicator dynamics for this phage-host system are:

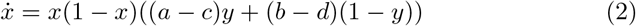

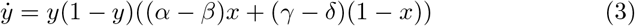

**Table 1:** Interpretations of parameter combinations in bimatrix game formulation of the Ferrojan Horse hypothesis.

| Parameter | Interpretation |
| --- | --- |
| $a - c$ | Relative fitness of cooperator host when infected by a Ferrojan virus |
| $b - d$ | Relative fitness of cooperator host when infected by a non-Ferrojan virus |
| $\alpha - \beta$ | Relative fitness of a Ferrojan virus infecting a cooperator host |
| $\gamma - \delta$ | Relative fitness of a Ferrojan virus infecting a defector host |

Where *x* represents the proportion of cooperator hosts and *y* represents the proportion of Ferrojan viruses. In Appendix 5.1, we derive this model from fitness calculations, solve the equilibria, and calculate their corresponding eigen-values (see Table 2).

**Table 2:** Eigenvalues of the Jacobian of bimatrix game theory model evaluated at all possible equilibria.

| $x^*$ | $y^*$ | $\lambda_1$ | $\lambda_2$ |
| --- | --- | --- | --- |
| 0 | 0 | $b - d$ | $\gamma - \delta$ |
| 0 | 1 | $a - c$ | $-(\gamma - \delta)$ |
| 1 | 0 | $-(b - d)$ | $\alpha - \beta$ |
| 1 | 1 | $-(a - c)$ | $-(\alpha - \beta)$ |
| $\frac{(\gamma - \delta)}{(\gamma - \delta) - (\alpha - \beta)}$ | $\frac{(b - d)}{(b - d) - (a - c)}$ | $\sqrt{\frac{(a - c)(b - d)(\alpha - \beta)(\gamma - \delta)}{((a - c) - (b - d))((\alpha - \beta) - (\gamma - \delta))}}$ | $-\sqrt{\frac{(a - c)(b - d)(\alpha - \beta)(\gamma - \delta)}{((a - c) - (b - d))((\alpha - \beta) - (\gamma - \delta))}}$ |

##### Mathematical Details

This system contains five equilibria, only one of which is a mixed strategy equilibrium, where both viral types and both host types stay at nonzero frequency. The mixed strategy equilibrium will only exist in *x* ∈ [0, 1] and *y* ∈ [0, 1] if the sign of (*a* − *c*) is the opposite of the sign of (*b* − *d*), and if the sign of (*α* − *β*) is the opposite of the sign of (*γ* − *δ*). The mixed strategy equilibrium can either be a saddle or the center of neutral periodic orbits depending on conditions described in Appendix 5.1.3. Due to symmetries in the eigenvalues, there can only be either one or two stable exterior fixed points as long as *a* − *c*, *b* − *d*, *α* − *β*, *γ* − *δ* are nonzero. When sgn(*a* − *c*) = sgn(*b* − *d*) or sgn(*γ* − *δ*) = sgn(*α* − *β*), there will only be one stable exterior fixed point. Otherwise, the system will have bistable exterior fixed points and a defined mixed strategy equilibrium.

##### Ecological Interpretation

In ecological terms, simultaneous existence of both viral and host types is only possible in this model when Ferrojan viruses have a fitness advantage infecting only one host type, and cooperator hosts have a fitness advantage when interacting with only one type of virus. If these conditions do not hold, then there will be a pure ESS for hosts and viruses. A pure ESS occurs when one host type and one virus type reach fixation and exclude invasion by other types. Next, we interpret this model in the context of two stable iron conditions: complete iron starvation and replete iron bioavailability.

#### 2.1.2 Stability Analysis of Bimatrix Model for Ferrojan Horse Hypothesis

First, we consider a scenario where the only bioavailable iron is iron ligated to a host’s siderophores. Under this assumption, producing siderophores will be necessary for host growth, so producers will always have a fitness advantage over non-producers. Using the expression for the payoff matrices in Eq 1, this means that the fitness of a cooperator host encountering a Ferrojan virus (*a*) will be higher than the defector host encountering the same virus type (*c*), and similar for non-Ferrojan viruses. This is equivalent to the conditions *a* − *c* > 0, *b* − *d* > 0. Using the signs of these two quantities and the information in Table 2, the feasible and stable equilibria will be either *x* = 1, *y* = 1 or *x* = 1, *y* = 0. The dynamics will approach one or the other depending on the signs of the relative fitness differences of viruses infecting cooperator hosts, i.e., whether *α* − *β* is positive or not. If *α* − *β* is positive, the only evolutionarily stable strategy (ESS) is host cooperation and the system will converge to one in which all viruses have iron tails. If *α* − *β* is negative, then the ESS for hosts remains the same but will be the opposite for viruses. If the two virus strategies have neutral fitness (*α* − *β* = 0), then any combination of viral strategies will be evolutionarily stable. A simulation of the *α* − *β* > 0 case is demonstrated in both phase space and time-evolution in Figure 2, left panel.

**Fig. 2:**
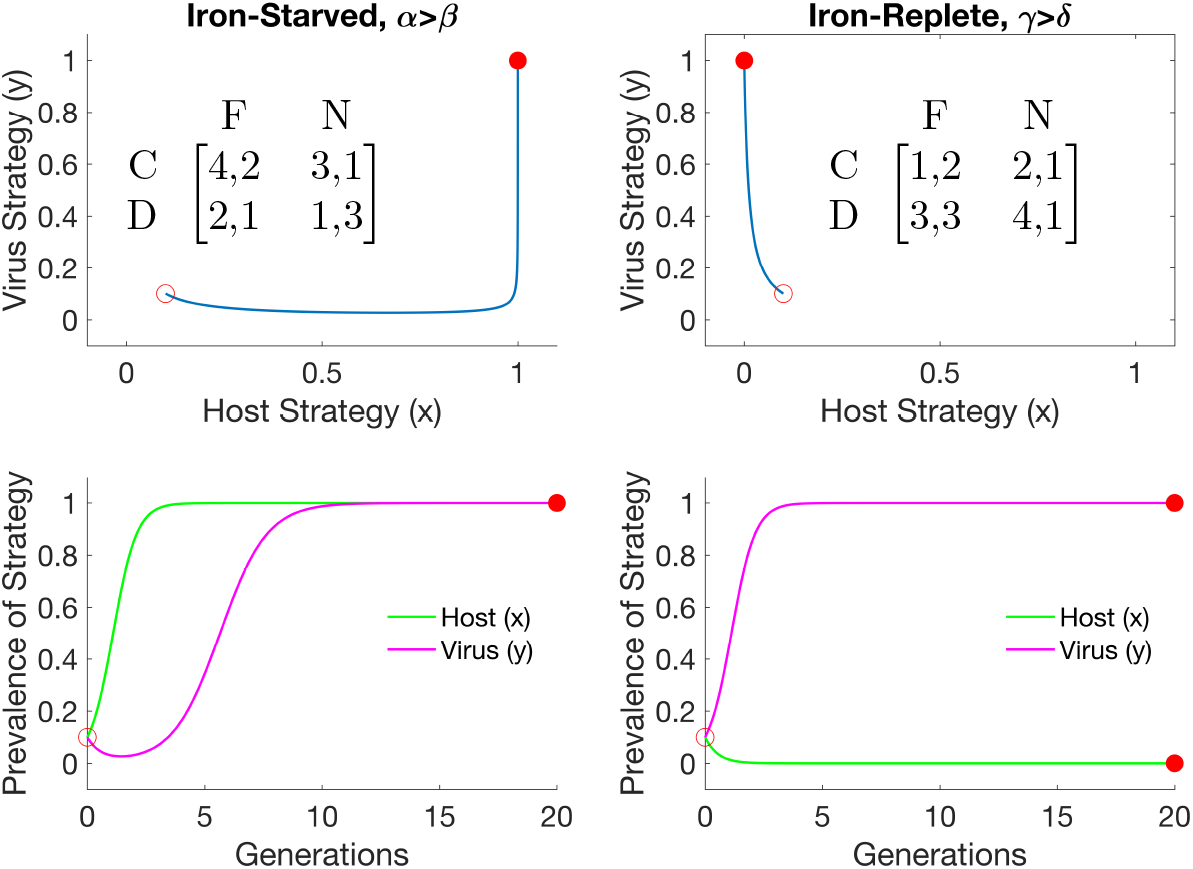
Phase diagrams and time evolutions of bimatrix nutrient-implicit model. Left panels show the dynamics of the model run in an iron-starved nutrient condition, with the payoff matrix shown in the phase plane. Right panels show the dynamics of the model run in the iron-replete nutrient condition and the corresponding payoff matrix. Open points indicate the initial conditions of each simulation, closed points indicate the equilibrium conditions.

Second, consider the scenario in which bioavailable iron is entirely replete. In this scenario, producing siderophores constitutes a fitness disadvantage. This is equivalent to the case *a* − *c* < 0, *b* − *d* < 0. As in the last scenario, the only ESS for hosts is *x* = 0 (no siderophore producers). The ESS for viruses will be no iron-tail incorporation if *γ* − *δ* < 0, all iron-tail incorporation if *γ* − *δ* > 0 (simulated in Figure 2 right panel), or any combination of strategies in the case *γ* − *δ* = 0. In both of these cases, we consider scenarios in which sgn(*a* − *c*) = sgn(*b* − *d*), such that there can only be one ESS for hosts (see Table 2).

### 2.2 Bimatrix Replicator Dynamics with Fluctuating Environment

Next, we consider the outcomes of the bimatrix game given a dynamical iron resource. To do so, we introduce a state variable, *n*, which refers to the relative bioavailability of iron for host growth. If *n* = 0, then bioavailable iron is depleted, and hosts must produce siderophores to ligate iron essential for growth. We emphasize that the environmental state *n* = 0 does not refer to a complete depletion of iron. Rather, it indicates that iron limitation has its strongest negative effect on host fitness in the absence of siderophore production. In contrast, if *n* = 1, then there is sufficient bioavailable iron such that hosts do not need siderophores for growth.

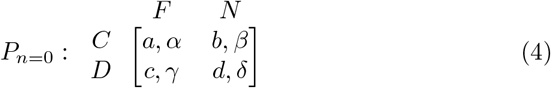

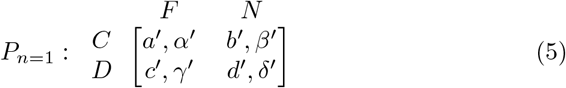

The payoffs at any given intermediate value of *n* is the linear interpolation of the *n* = 0 and *n* = 1 payoffs [47]. In this model, the payoffs for host and viral strategies will be a function of these two payoff matrices proportional to the current resource state. Specifically:

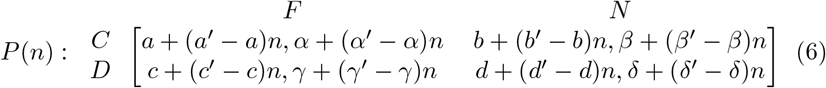

Although more general properties of the model will be analyzed, to investigate the biological question of viral impact on host siderophore evolution, we present some assumptions about model parameters below:

#### Host Payoff Conditions

We assume that during iron limitation (*n* = 0), producing siderophores is a higher-fitness strategy than not producing siderophores. Conversely, we also assume that under no iron stress (*n* = 1), not producing siderophores is a higher-fitness strategy. These assumptions are expressed in terms of model parameters in Table 3.

**Table 3:** Parameter constraints for host payoff matrices for environmental feedback-coupled bimatrix game.

| Condition | Interpretation |
| --- | --- |
| $a - c > 0$<br>$b - d > 0$ | Producing siderophores is advantageous<br>in iron-starved environments |
| $a' - c' < 0$<br>$b' - d' < 0$ | Producing siderophores is deleterious<br>in an iron-replete environments |

#### Virus Payoff Conditions

To interpret this model we also make assumptions about the fitnesses of viral strategies under differing iron limitation conditions. First, we assume that both host types express siderophore receptors, as well as the receptor for non-Ferrojan viruses, at comparable levels. Consequently, we assume that the fitness of a given viral strategy is independent of the host type the virus infects. While an argument could be made that possible differences in receptor expression between the two host types or increased viral-susceptibility due to the metabolic tax of siderophore production could lead to differential virus susceptibility, we leave results for these types of models to the Appendix. Under an iron-stressed condition, we assume that Ferrojan viruses have a fitness advantage due to the increased pressure to uptake iron via siderophore receptors. Conversely, in the iron-replete condition, we assume that siderophore-ligated iron outcompetes Ferrojan viruses for host uptake receptors, leading non-Ferrojan viruses to have a fitness advantage. Plaque experiments on cultures of *E. coli* grown with iron bound by the siderophore enterobactin show an inverse relationship between iron-enterobactin concentration and infection by phage H8, which uses siderophore-uptake receptors [36]. We describe these assumptions in terms of parameters values in Table 4.

**Table 4:** Parameter constraints for virus payoff matrices for environmental feedback-coupled bimatrix game.

| Condition | Interpretation |
| --- | --- |
| $\alpha - \beta > 0$<br>$\gamma - \delta > 0$ | Ferrojan viruses are more fit in iron-starved environments |
| $\alpha' - \beta' < 0$<br>$\gamma' - \delta' < 0$ | Ferrojan viruses are less fit when abundant siderophores compete for receptors |

#### Environmental Feedback Model

This replicator dynamical model is then coupled to a differential equation describing the relative change in environmental state. We assume that under no siderophore production, iron bioavailability decreases over time. We model this by including a logistic decay term in *ṅ*, following the functional form of [47]. We also assume that the proportion of the host population generating siderophores releases iron limitation at some rate *θ*_*x*_. This particular resource-model formulation is density independent. Recent study of unimatrix resource-coupled game theory by [45] suggests that using density dependent resource models may induce limit cycles in otherwise bistable dynamical regimes. Although their findings have not yet been extended to the bimatrix case, this leads us to believe that density-dependent resource models may be of future interest. Lastly for this model, we assume Ferrojan viruses reduce the environmental state by some rate *θ*_*y*_ by sequestering iron that otherwise might be available in dissolved lysate [33, 34].

Using these parameter and model attributes, we propose the following dynamical system:

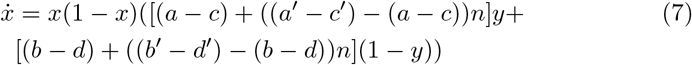

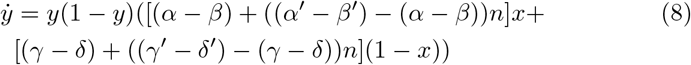

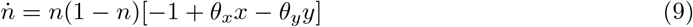

### 2.3 Classes of Model Behaviors

#### 2.3.1 Summary of Behavior Categories

Using the parameter conditions summarized in Table 3, we identify four classes of model behaviors for this system: (1) attraction to a single exterior fixed point, (2) neutral 2-dimensional orbits, (3) neutral 3-dimensional orbits, (4) attraction to a heteroclinic network (Figure 3). We interpret these dynamical behaviors as the following ecological situations: (1) resource crash, (2) dominating phenotypes, (3) phenotypic oscillations, (4) phenotypic jumping.

**Fig. 3:**
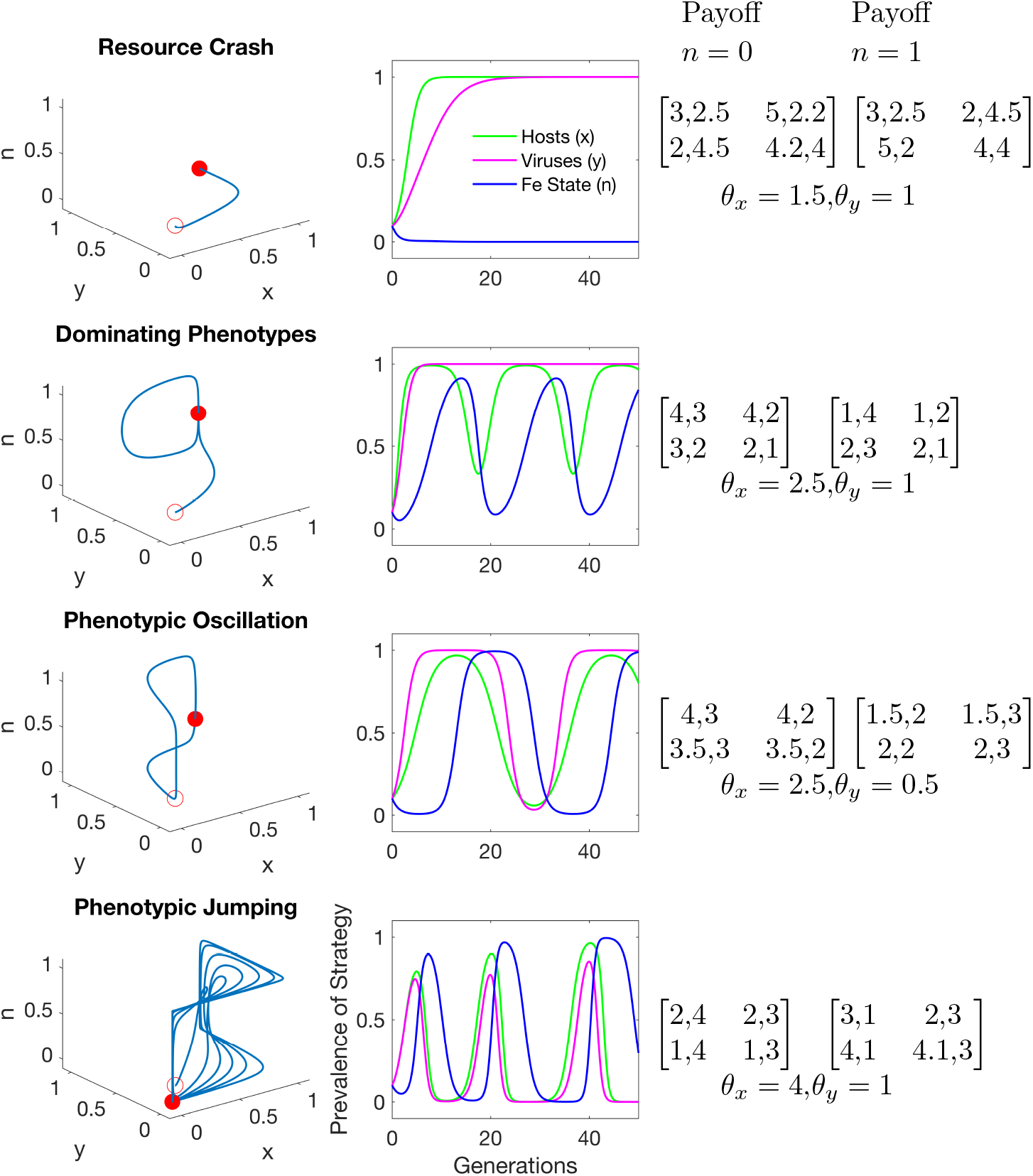
Phase diagrams and time evolutions for environmental feedback model exemplifying four classes of model behavior. Left panel is a phase diagram of the time evolution in center panels. Payoff matrices and resource-related parameters are listed in right panels. Each row corresponds to one of the classes of model behaviors. Open circle in phase diagram represents initial conditions, closed circle represents the state at the end of simulations (200 generations).

#### 2.3.2 Resource Crash

##### Mathematical Description

First, consider the case where host siderophore production is insufficient to generate a ‘replete’ iron state. This is equivalent to *θ*_*x*_ ≤ 1. Further, we assume that virus iron incorporation diverts iron from the available pool, i.e., *θ*_*y*_ > 0. With these parameters, *θ*_*x*_*x* − (1 + *θ*_*y*_*y*) < 0 for all *x, y* ∈ [0, 1], meaning *ṅ* will be negative in the entire state space. Any dynamics will then immediately go to the stable exterior fixed point where *n* = 0, *x* = 1 (because we enforce that cooperator hosts have an advantage in iron-starved conditions, see Table 3), and either *y* = 0 or *y* = 1 depending on which viral phenotype has higher average fitness infecting cooperator hosts.

##### Ecological Interpretation

Ecologically, the parameter *θ*_*x*_ represents the degree to which producing siderophores improves local iron bioavailability, *θ*_*y*_ represents iron incorporation into the tail fibers of Ferrojan viruses, and the −1 term indicates that iron limitation becomes more severe over time in the absence of siderophore production.Therefore, *θ*_*x*_ would only be less than 1 if a population of entirely cooperator hosts could not produce siderophores fast enough to improve local iron bioavailability, resulting in persistent iron starvation.

#### 2.3.3 Dominating Phenotypes

##### Mathematical Description

We define a ‘dominating strategy’ as a host or virus phenotype which has a fitness advantage in all environments. For example, Ferrojan viruses would be a dominating strategy for viruses if *α* − *β*, *γ* − *δ*, *α*′ − *β*′, *γ*′ − *δ*′ > 0. That is, if a Ferrojan virus has a fitness advantage over non-Ferrojan viruses infecting both cooperator and defector viruses in iron-replete or -deplete conditions. In the case of a dominating strategy for either viruses or hosts, one of the boundary planes of the state space will become attracting. Dynamics will always tend to that plane, and once that plane is reached, then the dynamics of the remaining two state variables will either be neutral orbits about an internal equilibrium, or that equilibrium will be a saddle and two of the corners of that plane will constitute bistable fixed points. We predict this behavior by solving for the constant of motion in terms of *x* and *n* for both the *y* = 0 and *y* = 1 planes (see Eq 31). We extend this finding and can identify a unique constant of motion for all fixed values of *y*, which will always exist given the ecological parameter constraints in Table 3 (see Eq 52). We can solve a similar Hamiltonian for the hosts in terms of *n* and *y*. The fixed point at the center of this Hamiltonian, however, is always unstable if there is a dominating viral phenotype, and will be unstable for some values of *x* ∈ [0, 1] otherwise (see Eq 60). We also note that these periodic oscillations can still occur even without a dominating strategy, as long as there is an unstable internal fixed point, explained in further detail in Appendix 5.3.

##### Ecological Interpretation

Using the assumptions in Table 3, cooperator hosts only have an advantage in iron limitation conditions and not in iron-replete conditions. Therefore, we assume no dominating strategy for hosts. Similarly, under the assumptions in Table 4, the Ferrojan strategy is only advantageous under iron-limitation and not in iron-replete conditions. However, in the case these assumptions are relaxed, we can assume, for example, that Ferrojan viruses always have a fitness advantage regardless of the extent of iron limitation. In this case our model predicts that the Ferrojan phenotype will rapidly reach fixation, after which the degree of iron limitation and proportion of cooperator hosts in the population will undergo neutrally stable periodic oscillations. This prediction contrasts with the bimatrix model with no environmental feedback, which predicts that a fixed viral phenotype cannot coexist with a host population of both cooperators and defectors.

#### 2.3.4 Population Oscillations

##### Mathematical Description

Neutral oscillations in environment quality as well as the frequency of phenotypes in both hosts and viruses can also occur in this model. Neutral 3D oscillations can only occur when there is a neutrally stable internal fixed point. Such a fixed point is feasible when there is some 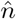 such that:

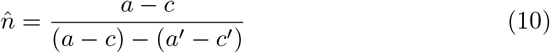

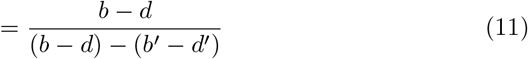

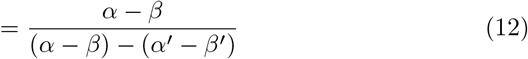

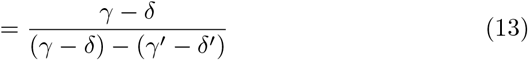

If 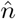 is non-negative in the range 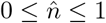, then when 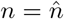, there will be no flow in the *x* or *y* directions. Further, *ṅ* = 0 when:

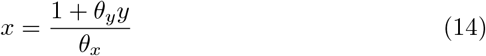

When 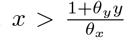, the flow will be towards *n* = 1, and towards *n* = 0 when 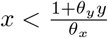. The intersection of the 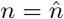 plane and the 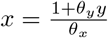 plane can be used to separate the state space into quadrants. In Appendix 5.2.5, we calculate the directions of flow with respect to these quadrants and demonstrate orbits. Additionally, we find under these conditions that all exterior fixed points are repelling while the interior fixed point is neutrally stable. The stability of fixed points are consistent with our finding of closed 3-dimensional periodic orbits.

##### Ecological Interpretation

While the oscillating-phenotypes outcome is interesting from a dynamical standpoint, due to a requirement of exact symmetries in the payoff matrices between nutrient states, we recognize this as a marginal case. Neutral oscillations in a similar nutrient-explicit bimatrix game theoretic model are also described in [16].

### 2.4 Characterizing Heteroclinic Networks

#### 2.4.1 Phenotypic Jumping

##### Mathematical Description

Using the payoff structure from Table 3, we find that all exterior fixed points will be unstable with either one or two positive eigenvalues if the resource crash is avoided, i.e., *θ*_*x*_ > 1 (see Appendix 5.3 for full demonstration). Heteroclinic cycles occur when the dynamics move in a repeating sequence from one unstable exterior fixed point to another, and are known to be common in replicator dynamical systems [23]. We identified heteroclinic cycles in this model using the characteristic matrix method developed by [18] (see Appendix 5.3).

Heteroclinic cycles can be characterized by the eigenvalues associated with each exterior fixed point in the cycle [6]. Heteroclinic networks occur when the heteroclinic cycles in a system are connected via positive transverse eigen-values [6]. In this model, heteroclinic cycles are guaranteed to exist when all three assumptions listed in Figure 5 are met. Additionally, heteroclinic cycles are guaranteed to form a heteroclinic network as long as any of the Ferrojan virus marginal fitnesses are positive. In terms of parameters, if any of *α* − *β*, *γ* − *δ*, *α*′ − *β*′, *γ*′ − *δ*′ are positive, there will be a heteroclinic network.

##### Ecological Interpretation

Heteroclinic cycles will appear as rapid transitions between the neighborhoods of external fixed points, where they linger for increasingly long times (see Figure 3 bottom center for example). In ecological terms, we can interpret this as rapid sweeps of either host or viral phenotype, or a rapid shift in iron limitation. Because each exterior fixed point is unstable, eventually small deviations will cause a new sweep to happen, and the system will ‘jump’ to a new state. The sequence of these ‘jumps’ is dependent on viral payoff structure, and multiple possible sequences can exist for one particular set of payoffs (see Figure 4).

**Fig. 4:**
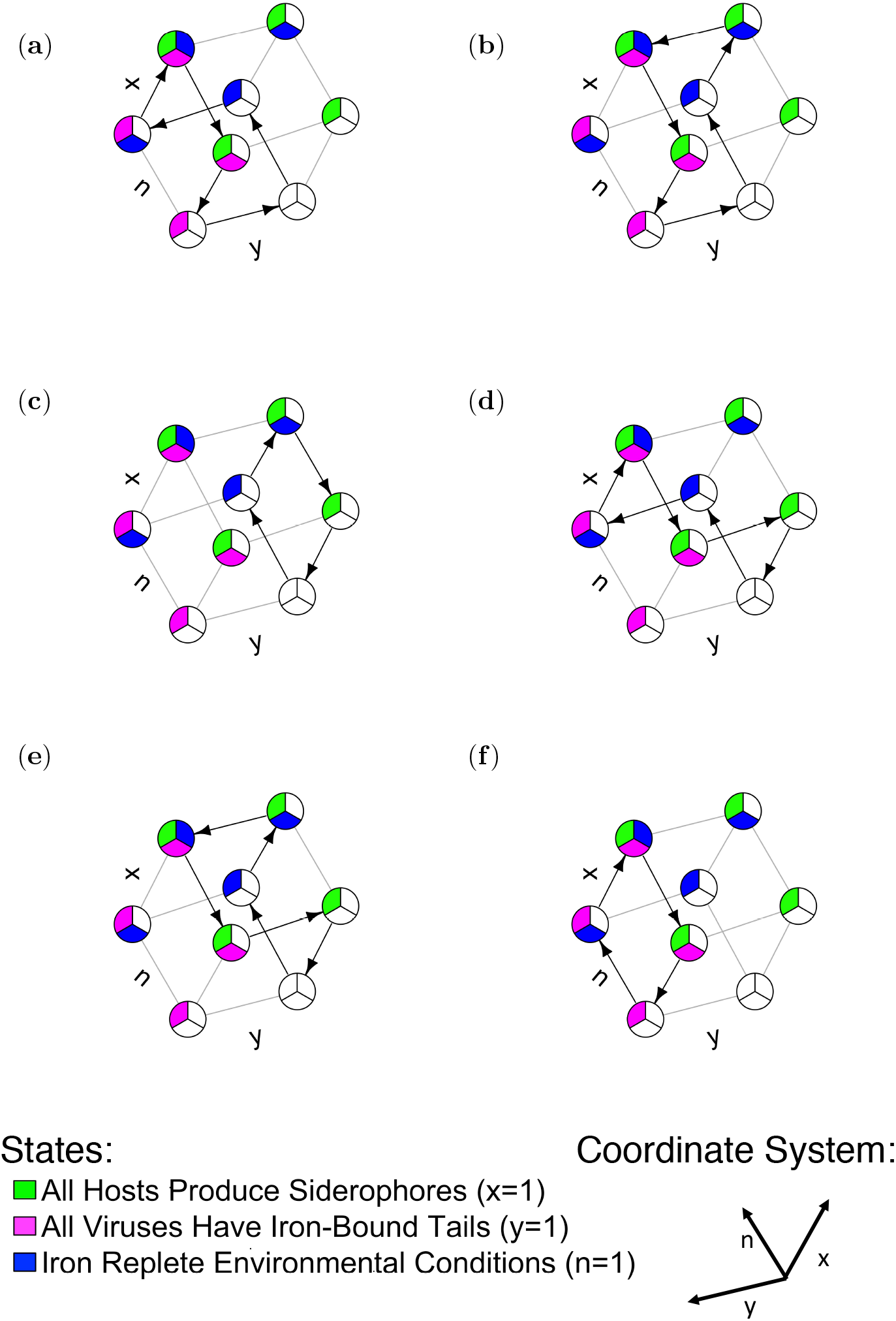
Diagram of heteroclinic network established using the same payoffs as the heteroclinic network panel of Figure 3. Nodes indicate exterior fixed points. Directed edges indicate trajectories of a heteroclinic cycle in the network. For this payoff structure, 6 heteroclinic cycles exist in the network, although at most only one cycle will be stable. The stable cycle observed in Figure 3 is cycle (**e**). Filled nodes correspond to the state indicated in the legend, unfilled nodes correspond to state 0.

**Fig. 5:**
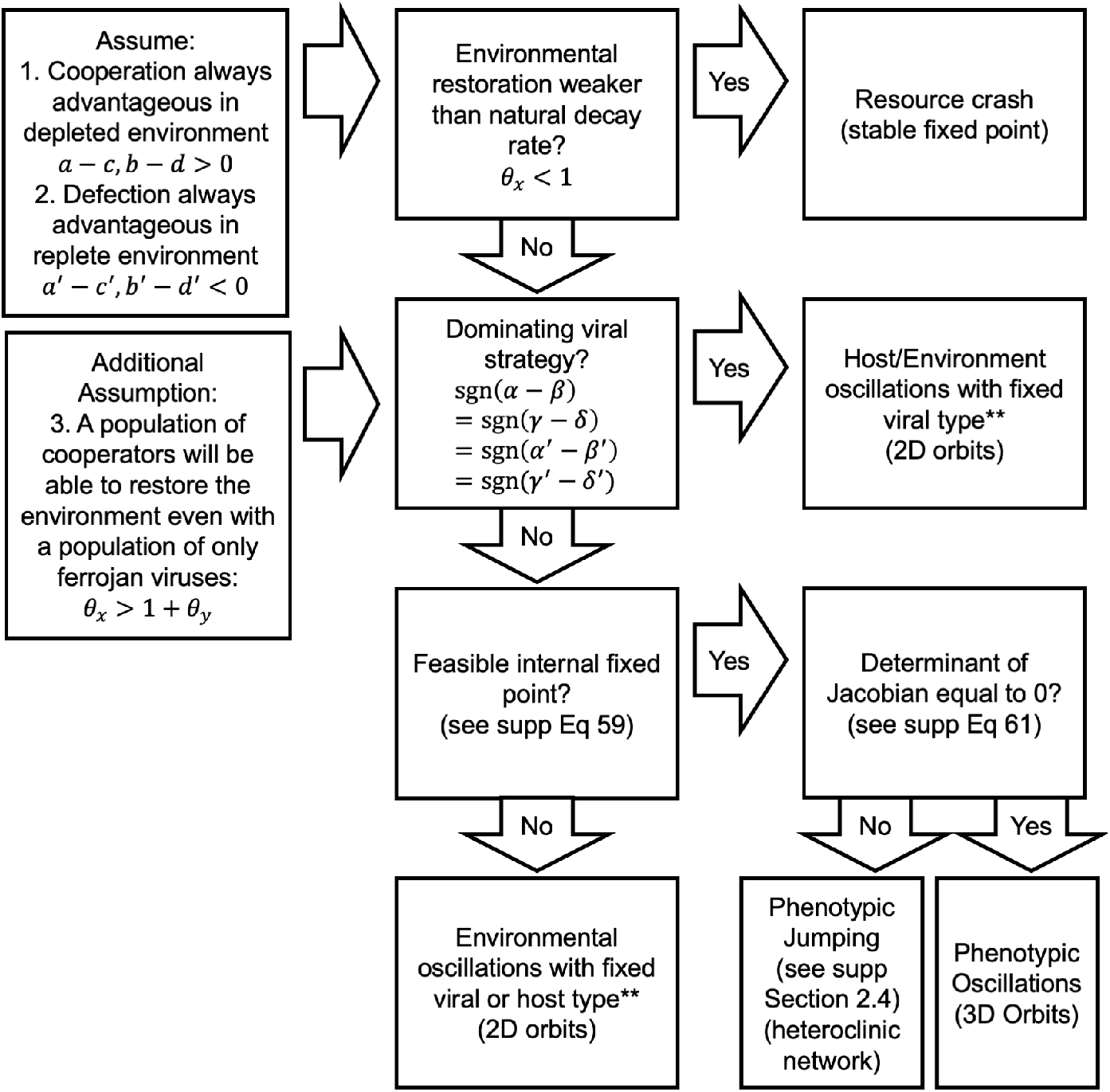
Flowchart summarizing parameter conditions which lead to particular model behaviors. We discuss the specifics for each of these behaviors in supplemental information. ** Denotes that 2D orbits broadly refer to Hamiltonian systems, where in supplemental section 2.2 we delineate parameter conditions for neutral orbits about an internal fixed point versus bistable exterior fixed points. Briefly, if assumption 3 is met, then neutral oscillations will occur for dynamics on the *y* = 0, 1 planes, neutral oscillations cannot occur for the *x* = 0 plane, but can occur for the *x* = 1 plane.

**Fig. 6:**
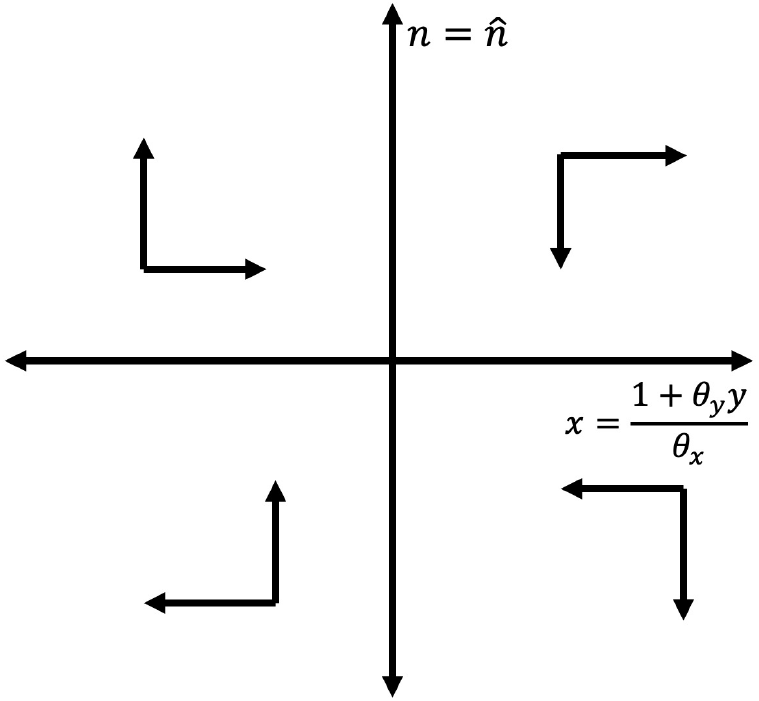
Directions of flow for 3D neutral orbits. Vertical axis indicates the 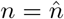 plane and horizontal axis indicates 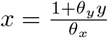 plane.

**Fig. 7:**
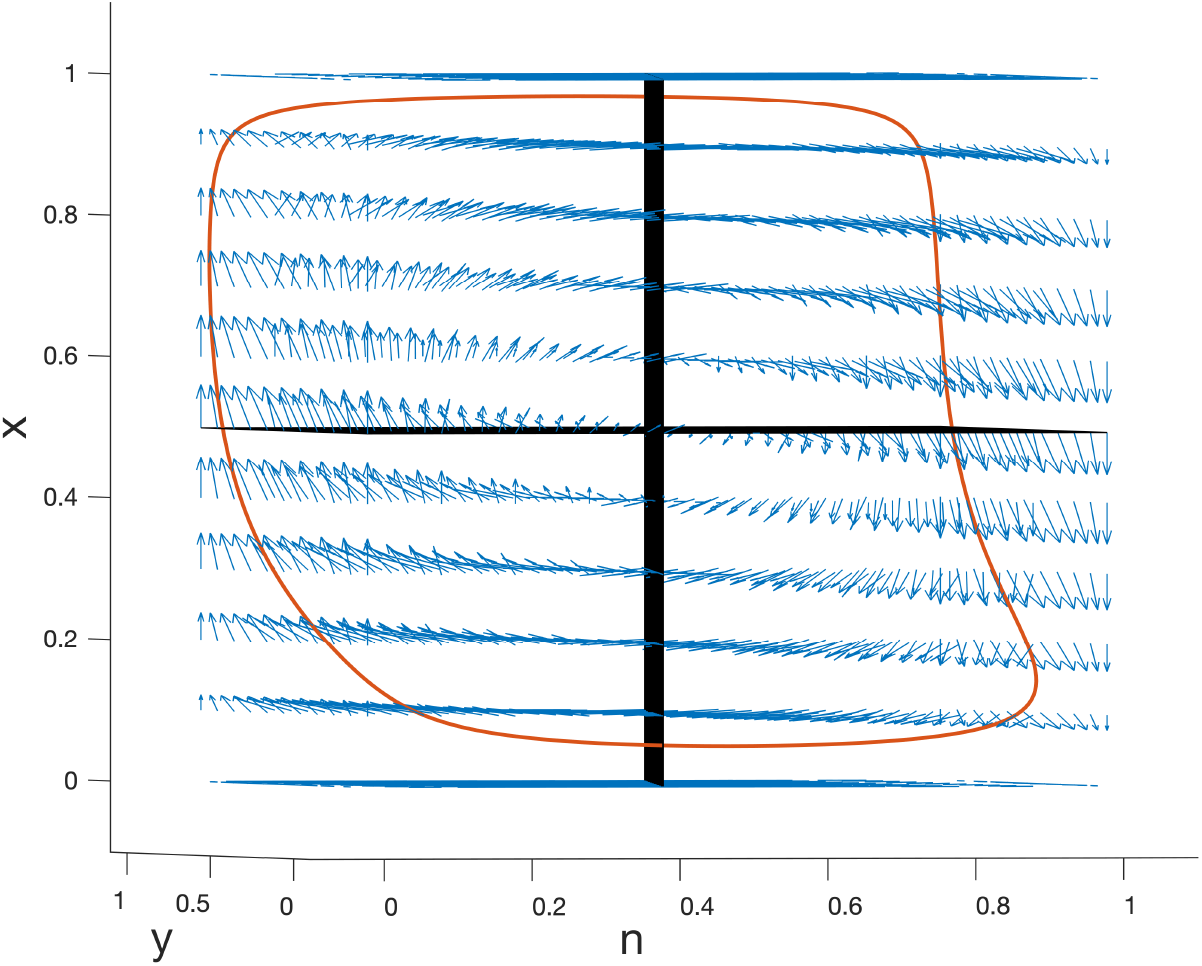
Phase portrait of flow for 3D neutral orbits. Black lines indicate null-clines for *ṅ* and *ẋ*, blue arrows indicate flow, and red line indicates sample orbit trajectory.

**Fig. 8:**
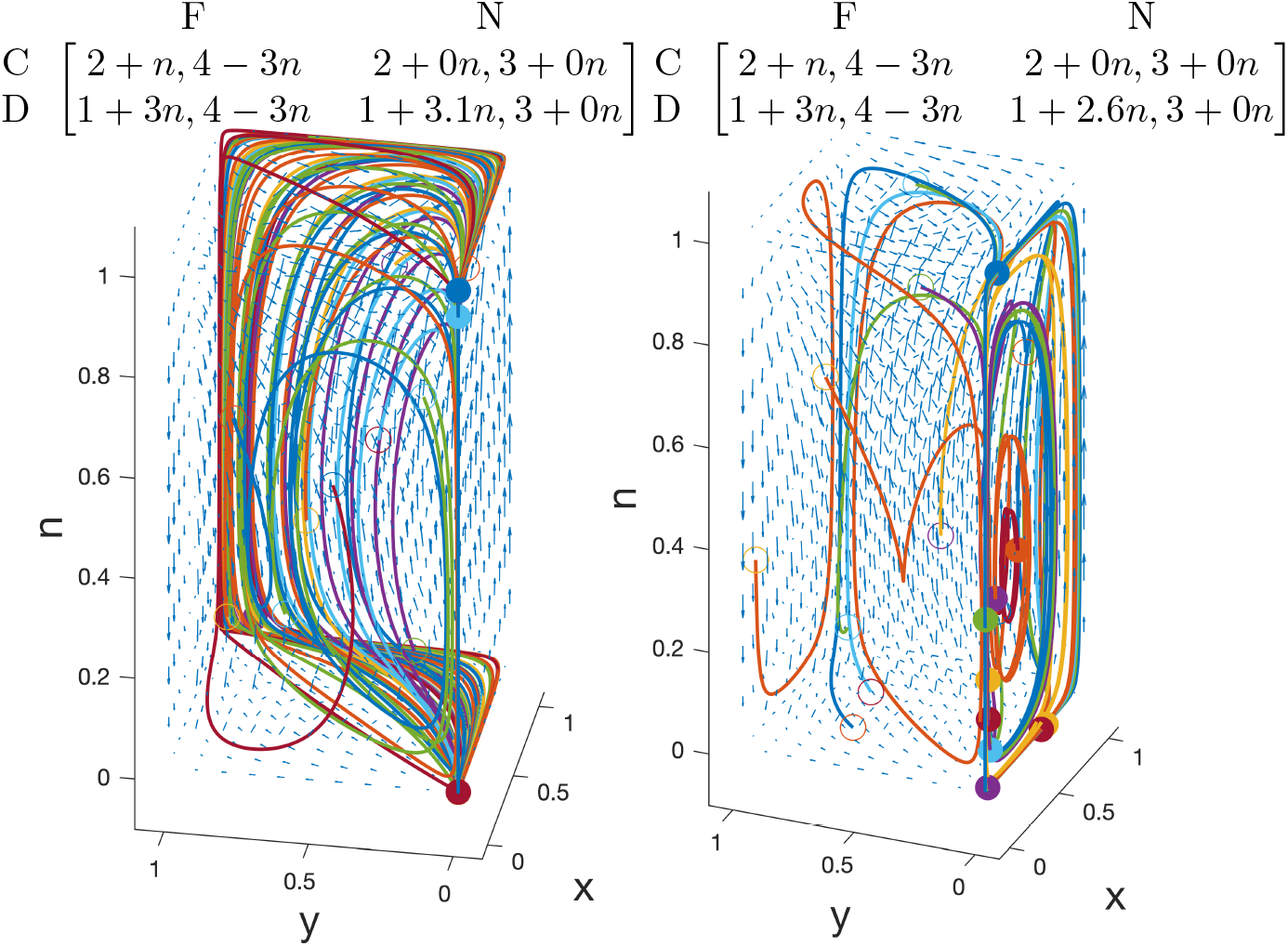
Comparison of dynamical outcomes for models with the same heteroclinic networks, but where one model has an unstable interior fixed point (left) and the other has no defined interior fixed point (right). Ten trajectories with different initial conditions are displayed. Colors indicate sample trajectories, arrows indicate flow fields. Open points are initial conditions closed points are conditions at 200 generations (the end of the simulation).

To illustrate the interpretation of heteroclinic cycles, we provide two concrete examples of heteroclinic cycles belonging to the heteroclinic network demonstrated in Figure 3 as represented by panels (**a**) and (**e**) in Figure 4. In cycle (**a**), consider a system that starts with an initial condition under extreme iron stress with no siderophore producing hosts and no Ferrojan viruses (the completely unfilled node). While the concentration of siderophore-bound iron is still low, Ferrojan viruses have a fitness advantage over non-Ferrojan viruses and sweep the population. Once this occurs, siderophore-producing hosts sweep the host population, after which iron bioavailability improves. Upon improved iron conditions, non-producer hosts sweep the host population, and competition with siderophores for uptake causes Ferrojan viruses to be outcompeted by non-Ferrojan viruses. However, with a population of all non-producer hosts, iron stress is reintroduced and the cycle repeats. Cycle (**e**) (the attracting cycle in this system, as shown in Figure 3) tells a slightly different eco-evolutionary story. Here, start again in a condition under extreme iron stress with no Ferrojan viruses or siderophore-producing hosts. First, siderophore-producing hosts sweep the host population. Then, Ferrojan viruses sweep the viral population while iron stress is still high. Siderophore-producing hosts increase iron bioavailability and relieve iron stress, after which Ferrojan viruses are out-competed by non-Ferrojan viruses. Then, under low iron stress, non-producer hosts sweep producers, after which iron bioavailability declines and the cycle restarts. Using the intuition that cycles with the largest ratio of contracting-to-expanding eigenvalues will ultimately be attracting neatly dovetails with our ecological interpretations, as the eigenvalues in the *x* and *y* directions associated with each exterior fixed point simplify to the relative fitnesses of host and virus phenotypes at the local environmental state (see Appendix 5.3).

## 3 Discussion

Viral infection [15] and demand for iron [28] are both important aspects of marine microbial ecology. The Ferrojan Horse Hypothesis offers a potential mechanistic linkage between these two factors. We use a bimatrix game theoretic model to assess the eco-evolutionary impacts of the Ferrojan Horse hypothesis on host siderophore use. In a nutrient-implicit model, we find that viral infection does not impact the global evolutionary dynamics of the hosts. Instead, the host phenotype which performs better in a given iron condition always reaches fixation, regardless of its susceptibility to viral infection. We then coupled this model to an explicit environmental feedback. Under varying environmental states, the host and environmental eco-evolutionary dynamics of this model strongly depend on viral strategies. If a single viral phenotype has a fitness advantage under all conditions, the model predicts oscillatory dynamics for both host phenotype and environmental state. Under different viral fitness conditions, dynamics attract to a heteroclinic network, which implies intermittent rapid switching between extreme host phenotype, virus phenotype, and environmental states. All of these diverge from the expectation under a constant-environmental model, which predicts a single, pure host ESS.

This model contains some simplifying assumptions. Notably the bimatrix game formulation considers frequency-dependent but not density-dependent effects. As a consequence, the absolute concentration of iron is not modeled. In related work considering games with environmental feedback, Tilman et al [45] show how alternative, density-explicit environmental state models can alter system dynamics by inducing limit cycles. Extending the present model to a density-dependent context is a priority for future research; it is possible that emergent limit cycles may exist for different resource models capitulated in *ṅ*. In addition, the present model does not take into account the effects of spatial heterogeneity on dynamics. Previous research in ecological games shows that local interactions can have qualitative impacts on model behavior, such as stabilizing coexistence or inducing limit cycles [25, 17]. Studies in aquatic environments also show the importance of spatial heterogeneity for microbial community composition [43], gene expression [40], and viral populations [42]. In addition to spatial heterogeneity, this model framework can also be adapted to study heterogeneity in host receptor expression levels [31]. This modeling framework can continue to study whether expressing siderophore uptake receptors at low levels is more or less evolutionarily advantageous in terms of the viral susceptibility/iron uptake tradeoff, and to what extent local environmental conditions may impact siderophore production and uptake in an evolutionary sense. Although genome regulation under iron limitation for marine cyanobacteria has largely focused on the down-regulation of cellular machinery with high iron content [29] or intracellular recycling of iron [39], it is also possible siderophore-uptake receptor expression may be regulated by environmental conditions.

In closing, via our resource-coupled bimatrix replicator dynamical modeling approach, we find viral infection can have a substantive impact on host adaptation to iron scarcity under variable resource conditions. Further, viral phenotypes as well as their impacts on host adaptation propagate to affect the dynamics of the resource itself. Our model suggests that the Ferrojan Horse hypothesis warrants further investigation in marine environments, as viruses may have yet to be understood impacts on iron dynamics on microbial communities [13]. Understanding viral impacts on marine microbial communities is necessary to disentangle bottom-up and top-down drivers of global primary production. Incorporating viruses into models of microbe-iron feedbacks will also help improve predictions of iron amendments and their consequences for carbon drawdown in an ecosystem context.

## 4 Materials and Methods

### 4.1 Code Availability

All numerical integrations of ODEs were carried out via the 4th order Runge-Kutta method as implemented by function ode45 in MATLAB v 2017a. Heteroclinic networks were visualized using R v 3.5.2’s [35] igraph v 1.2.2 [12] package. Scripts are available at at https://zenodo.org/badge/latestdoi/191977232.

## Acknowledgements

The authors would like to thank Stephen Abedon, David Talmy, Mya Breitbart, and Rachel Kuske for their helpful input on manuscript drafts. The authors also thank Stephen J Beckett for code review, and the Simons Collaboration on Ocean Processes and Ecology (SCOPE) community for their support. This research was funded by the Simons Foundation (SCOPE Award ID 329018).

## 5 Appendix

### 5.1 No feedback model

#### 5.1.1 Derivation from fitnesses

To build the bimatrix replicator model with no environmental feedback, begin with the payoff matrix:

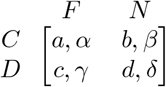

The average fitnesses (*r*) of each strategy for both hosts and viruses can be determined by the payoff matrix and the frequencies of each strategy. Denote the frequency of cooperator hosts as *x* and the frequency of Ferrojan viruses as *y*, where the frequency of defector hosts are 1 − *x*, and of non-Ferrojan viruses are 1 − *y*. The fitness values are:

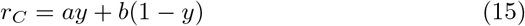

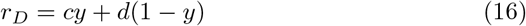

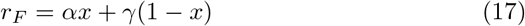

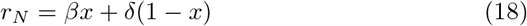

Where the subscripts *C, D, F, N* denote cooperator hosts, defector hosts, Ferrojan viruses, and non-Ferrojan viruses, respectively. The change in frequency of a strategy increases proportionally to the difference between that strategy’s fitness and the entire population’s average fitness. The dynamics for the prevalence of host and viral strategies becomes:

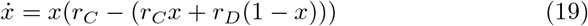

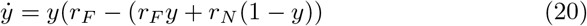

Which simplifies to:

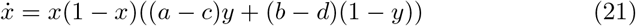

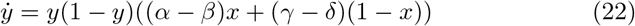

#### 5.1.2 Identifying fixed points

The nullclines of Eq 21 and 22 are:

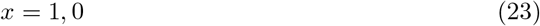

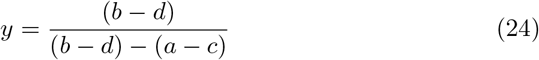

*y* nullclines:

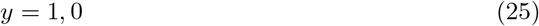

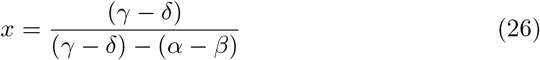

with fixed points listed in Table 5.

**Table 5:**
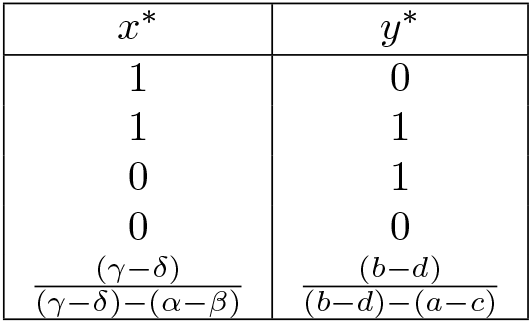
Fixed points of bimatrix game

| $x^*$ | $y^*$ |
| --- | --- |
| 1 | 0 |
| 1 | 1 |
| 0 | 1 |
| 0 | 0 |
| $\frac{(\gamma-\delta)}{(\gamma-\delta)-(\alpha-\beta)}$ | $\frac{(b-d)}{(b-d)-(a-c)}$ |

#### 5.1.3 Stability Analysis

We linearize the system about each of the fixed points to determine their local stability. The Jacobian is:

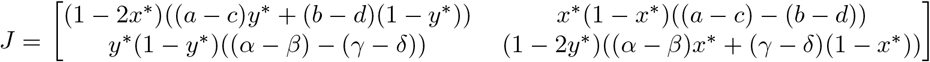

For the external fixed points, the off-diagonals of the Jacobian simplify to 0, so the eigenvalues correspond to the diagonal elements. The eigenvalues for each boundary fixed point are listed in Table 6.

**Table 6:** External eigenvalues for bimatrix game.

| $x^*$ | $y^*$ | $\lambda_1$ | $\lambda_2$ |
| --- | --- | --- | --- |
| 1 | 0 | $-(b-d)$ | $\alpha - \beta$ |
| 1 | 1 | $-(a-c)$ | $-(\alpha - \beta)$ |
| 0 | 1 | $a - c$ | $-(\gamma - \delta)$ |
| 0 | 0 | $b - d$ | $\gamma - \delta$ |

For the interior fixed point, we solve:

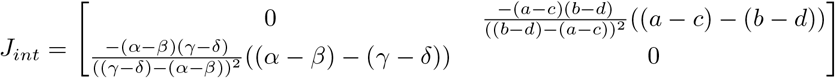

For this fixed point, the eigenvalues are:

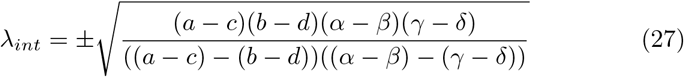

If the quantity inside this radical is positive, the internal equilibrium will be a saddle, but if it is negative, the eigenvalues will be imaginary and stability must be further investigated. For this case, we identify closed neutral orbits by solving the Hamiltonian of this system. To do this, we begin with 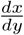:

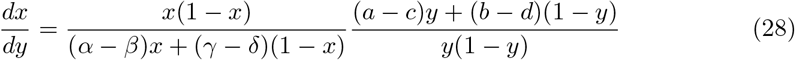

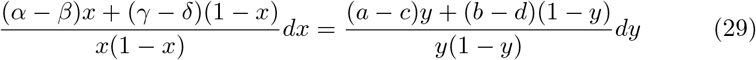

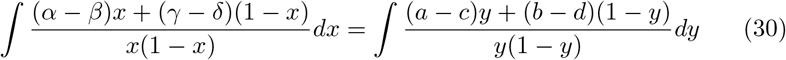

Solving with partial fractions yields

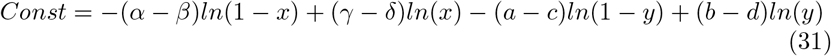

We confirm this Hamiltonian is time-independent, in that 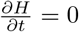

#### 5.1.4 Conditions for Neutral Stability

First, for there to be an internal fixed point with neutral stability, that fixed point must be feasible, i.e., reside in *x, y* ∈ [0, 1]. Using the expressions for *x*^*^, *y*^*^ in Table **??**, the following conditions are necessary for the fixed points to be feasible:

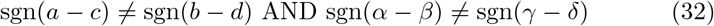

When this condition is imposed, the numerator in the radical for the expression the eigenvalues associated with this fixed point is necessarily nonnegative. Therefore, for neither eigenvalue to have a positive real part, the denominator must be negative, so either

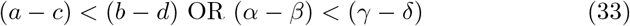

We conclude that this system will exhibit neutral periodic oscillations along a constant energy surface if both 32 and 33 are true.

### 5.2 Feedback model

#### 5.2.1 Derivation from fitnesses

For the feedback model, we construct two payoff matrices, one to represent a replete environment, and one to represent a depleted environment. For intermediate environmental conditions, we assume the payoff matrix is a linear interpolation of these two matrices. Therefore, the payoff matrix becomes a function of environmental condition, which is expressed as:

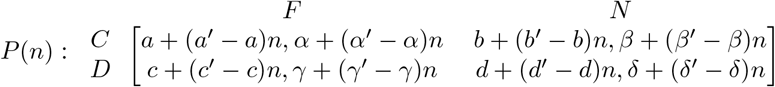

Then, we use the same derivation as in the no feedback model to define dynamics for *x* and *y* in terms of *n*. These are the new fitnesses:

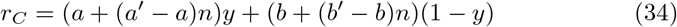

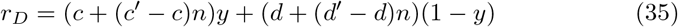

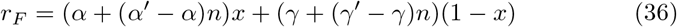

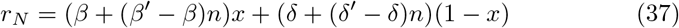

Using these fitnesses gives the following dynamics:

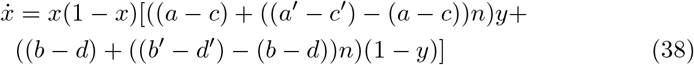

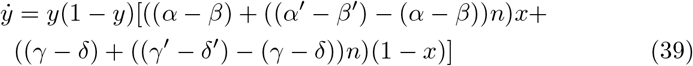

Now, dynamics must be established for *n*. We use a logistic decay term to restrict *n* to [0, 1]. Then, we increase *n* proportional to the fraction of cooperator hosts and decrease *n* relative to the fraction of Ferrojan viruses. This results in the following dynamics for n:

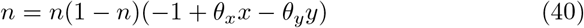

We assume that cooperator hosts will always have a fitness advantage regardless of which type of virus they encounter when there is no bioavailable iron. We further assume that defector hosts will always have a fitness advantage in an environment replete with iron. We represent these effects with the parameter constraints in Table 7.

**Table 7:** Parameter constraints for payoff matrices for environmental feedback-coupled bimatrix game.

| Condition | Interpretation |
| --- | --- |
| $a - c > 0$<br>$b - d > 0$ | Producing siderophores is always advantageous<br>in iron-starved environments |
| $a' - c' < 0$<br>$b' - d' < 0$ | Producing siderophores is always deleterious<br>in iron-replete environments |

#### 5.2.2 Dynamical Behaviors

#### 5.2.3 Resource Crash Demonstration

Here we demonstrate the resource crash dynamical outcome. Start with the environmental dynamics:

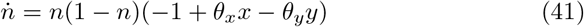

Because *n* ∈ [0, 1], both the *n* and 1 − *n* terms will always be positive. When *θ*_*x*_ ≤ 1, *θ*_*x*_*x* ≤ 1 ∀*x* ∈ [0, 1]. Therefore, the (−1 + *θ*_*x*_*x* − *θ*_*y*_*y*) term will always be negative, and *ṅ* will always be negative, meaning dynamics will always go to *n* = 0. In this model, when *n* = 0, cooperation is a dominant strategy for hosts, and so dynamics will also go to *x* = 1. The edge *x* = 1, *n* = 0 is flow invariant. Trajectories go toward *y* = 1 if Ferrojan viruses have a fitness advantage on cooperator hosts in a scarce environment (i.e., *α* − β > 0), or *y* = 0 in the opposite case.

#### 5.2.4 Solving Hamiltonians in 2D

We find a Hamiltonian in this system similar to the no-feedback game. We demonstrate these for any fixed value of *y* or *x* on an *x* − *n* or *y* − *n* plane, respectively. First, select some fixed value of *ŷ*. Then calculate 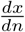:

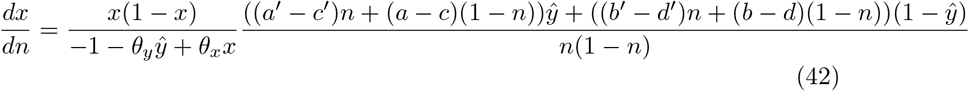

We can integrate this function using partial fractions. The result is:

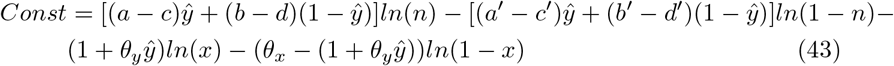

We confirm that 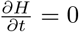. Now analyze the stability of the interior fixed point of this plane. We calculate this fixed point in terms of *ŷ*:

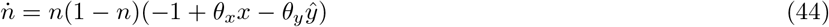

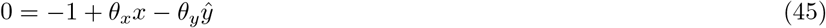

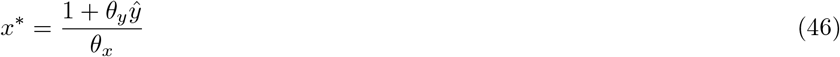

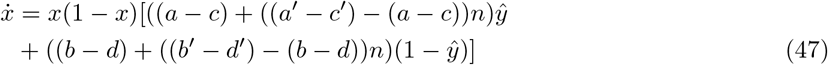

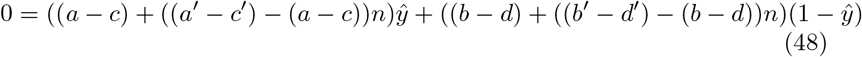

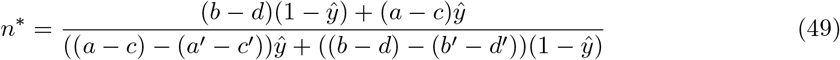

If *y* is held constant at *ŷ*, we can evaluate a Jacobian using *ẋ* and *ṅ* at this fixed point and determine its eigenvalues. The expression for the eigenvalues becomes:

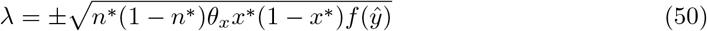

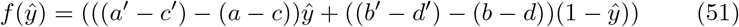

As long as *x*^*^, *n*^*^ ∈ [0, 1], one of the eigenvalues will have a positive real part unless:

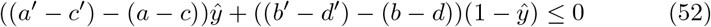

The payoff structure we enforce for this model includes (*a − c*) > 0 > (*a*′ − *c*′) and (*b* − *d*) > 0 > (*b*′ − *d*′). Therefore, we can conclude for all *ŷ* ∈ [0, 1] the *x* and *n* dynamics can be described as neutral periodic oscillations about a closed surface. This conclusion is important for choices of *ŷ* where *ẏ* = 0, i.e., *ŷ* = 0, 1. For other selections of *ŷ*, the dynamics of *y* will prevent the system from staying on a neutral 2D *x* − *n* orbit. We also note that the equilibrium coordinate *x*^*^ will be in the domain for all selections of *ŷ* as long as *θ*_*x*_ > 1 +*θ*_*y*_. In biological terms, there can be neutral oscillations between host types and environmental iron conditions for any fixed frequency of Ferrojan viruses if a population of only cooperator hosts can still improve iron conditions with a population of entirely Ferrojan viruses.

In a similar manner to the orbits we just solved, we can solve orbits for some fixed value of 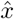 on the *y* − *n* plane. We find:

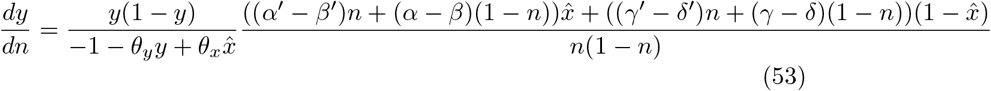

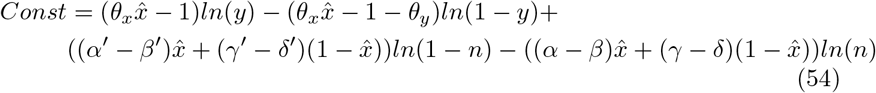

Which is, again, a time-independent Hamiltonian with the internal fixed point:

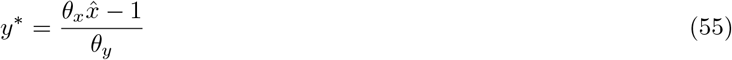

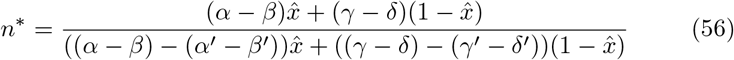

with eigenvalues:

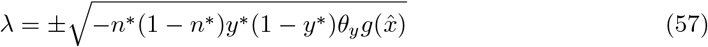

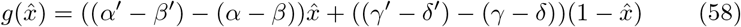

For this fixed point to be neutrally stable, the expression 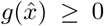. With *y*^*^, *n*^*^ ∈ [0, 1], we write a condition for 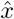 which is equivalent to 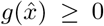.

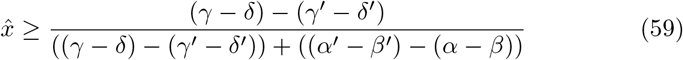

When this inequality is true, the dynamics of *y* − *n* for any 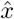 will be neutral periodic orbits. Further, we can examine this condition in more detail. First, in the case where sgn((*γ* − *δ*) − (*γ*′ − *δ*′)) = sgn((*α* − *β*) − (*α*′ − *β*′)), the condition will be equivalent to 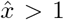, which means there will be no neutral orbits for any selection of 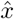 in the domain of *x*. If either sgn((*γ* − *δ*) − (*γ*′ − *δ*′)) or sgn((*α* − *β*) (*α*′ − *β*′)) is zero, the fixed point for *n*^*^ will be undefined, so there will be no neutral oscillations. Finally, if sgn((*γ* − *δ*) (*γ*′ − *δ*′)) ≠ sgn((*α* − *β*) (*α*′ − *β*′)), there will some values of 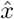 for which neutral oscillations are possible. We can additionally constrain this value, because the *y* coordinate of the fixed point must also be in the domain of *y*. Leveraging the expression for *y*^*^, we find:

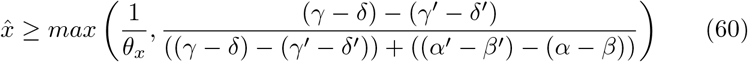

For 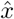 below these greater of these two values, neutral *y* − *n* orbits along a conserved surface are impossible. This condition is different from the condition for *x* − *n* orbits, which we find to be possible for any selection of *ŷ* given the parameter constraints in Table 7. However, because this range does not include 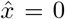, the only selection of 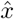 for which neutral periodic oscillations are an equilibrium is 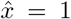. This means there can only be oscillations in Ferrojan/non-Ferrojan virus phenotypes for a host population of only cooperators.

Finally, we can extend the Hamiltonian solved in 5.1.2 for the 3D case for a constant value 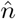. The results of this analysis are analogous, the Hamiltonian is:

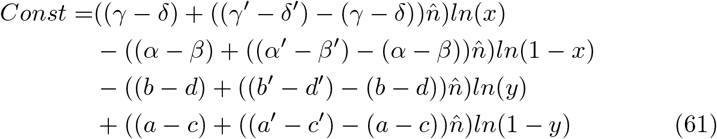

We can then evaluate the interior fixed point and its local stability and find:

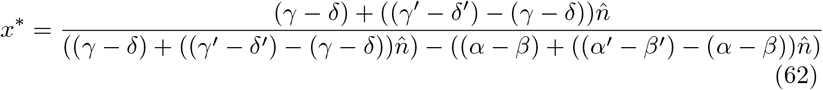

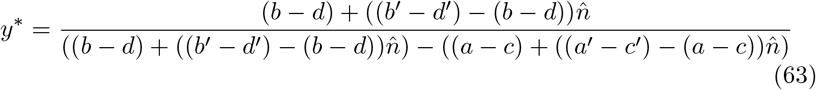

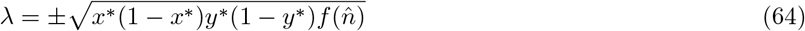

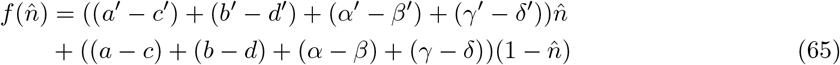

As long as 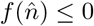, neutral periodic orbits will occur in the *xy* plane for any constant value of 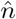. We can rewrite this condition as:

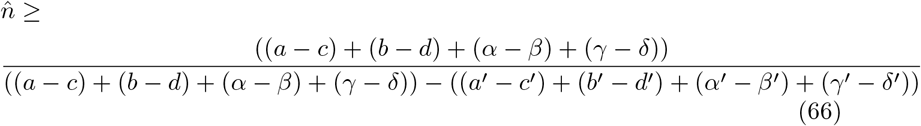

#### 5.2.5 3D Neutral Orbits

#### 5.2.6 Identifying Internal Equilibrium

We derive the nullclines of the entire model to understand the interior fixed points.

*x* nullclines:

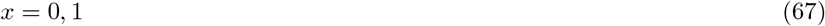

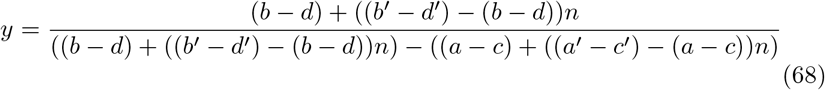

*y* nullclines:

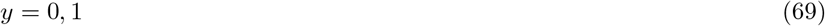

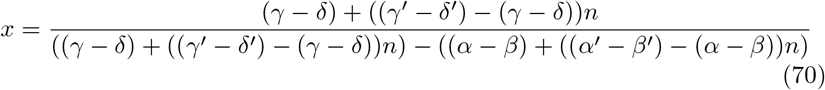

*n* nullclines:

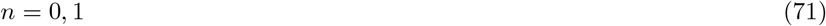

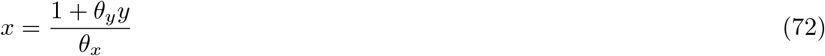

Solving the intersection of the nullclines, the interior fixed points can be described by the solutions to this quadratic equation of n:

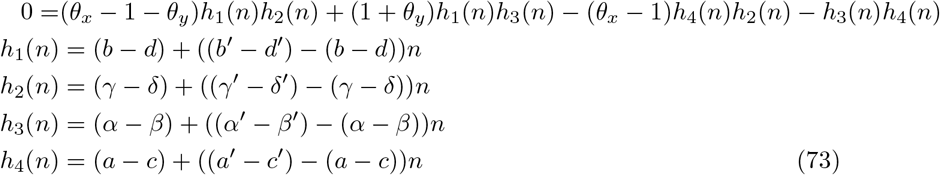

The system will have either 0,1, or 2 interior fixed points for any given payoff structure. Next, we determine the Jacobian evaluated at these fixed points:

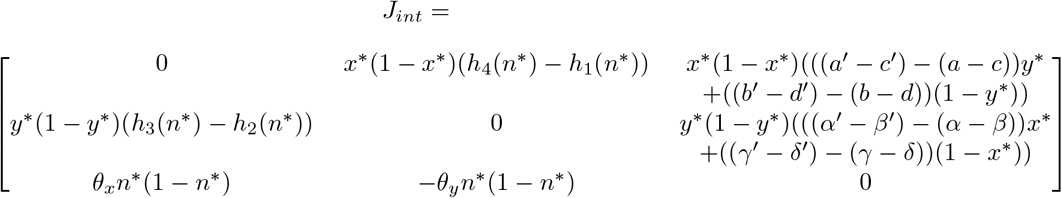

We evaluate the trace and determinant of this matrix:

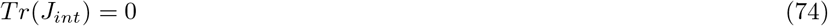

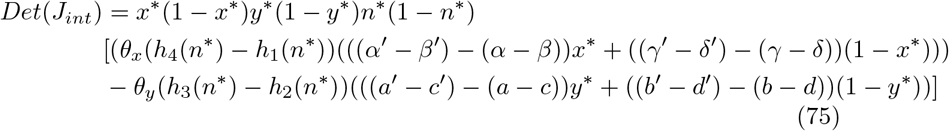

Since the trace of the Jacobian is equal to zero, the sum of the eigenvalues must be equal to zero. Using the complex conjugate root theorem, there are two possible combinations of eigenvalues. Either the eigenvalues will be a complex conjugate pair and one real number, or three real numbers. For three real numbers to sum to zero, they must either all be zero or there must be at least one positive and one negative number. If the eigenvalues are one real number and one conjugate pair, to sum to zero either they must all have a zero real part, or the sign of the real part of the complex eigenvalues will be opposite the sign of the real eigenvalue. If any of the eigenvalues have a positive real part, then the fixed point will be unstable. We therefore conclude that the interior fixed point will be unstable unless all eigenvalues have zero real part. If all eigenvalues have zero real part, then their product must be zero. The product of the eigenvalues of a matrix are equal to its determinant. Therefore, we find neutrally stable fixed points will only exist when the expression for the determinant of the Jacobian in Eq 75 is equal to zero. Otherwise, the interior fixed points will be unstable. We can summarize the parameter conditions that determine the four classes of dynamical outcomes presented here as a flowchart in Figure 5.

#### 5.2.7 Demonstration of orbital flow about null-planes

Under some parameterizations, it is possible that both *x* and *y* can be invariant for some particular value of *n*, 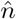. This occurs when:

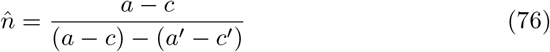

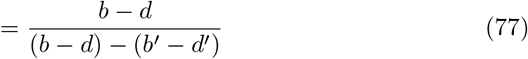

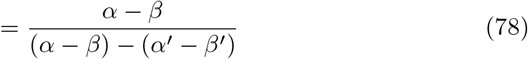

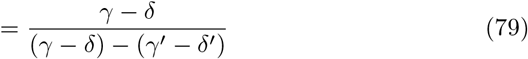

If 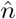 exists, when 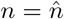, there will be no flow in the *x* or *y* directions. There is a corresponding plane describing when *ṅ* = 0, which has the following definition:

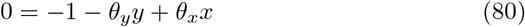

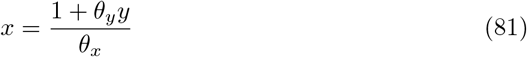

Now consider *ṅ* for values of *x*. If 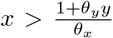 Here *ṅ* > 0. For values of *x* such that 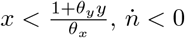. Substituting the equality for the *n* nullcline into *ẋ*:

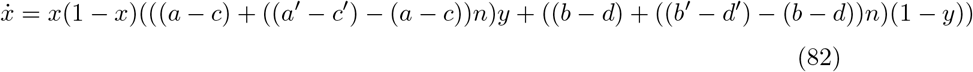

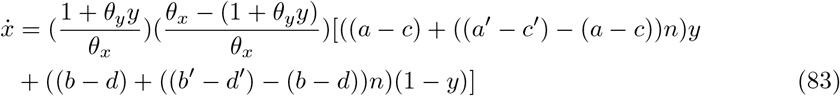

The first term in this product will be positive for all *y*. The second term will be positive for all *y* as long as *θ*_*x*_ > 1 + *θ*_*y*_. Under that condition the sign of *ẋ* will only depend on the third term. When 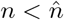, this term will be positive for all *y*, and negative for all *y* when 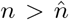. We represent the directions of the flow diagrammatically in Figure 6. This field suggests clockwise orbits such as observed when the model is simulated under the parameter conditions described in the main text (see Figure 7).

### 5.3 Heteroclinic Networks

#### 5.3.1 Stability Analysis of Exterior Fixed Points

We also analyze the stability of the exterior fixed points. For any of the exterior fixed points, the Jacobian becomes diagonal, and so the eigenvalues can be read off of the matrix. The eigenvalues for each exterior fixed point are summarized in Table 8. All of these fixed points have at least one positive eigenvalue using parameter constraints that avoid resource crash dynamics. All fixed points also have at least one negative eigenvalue if *θ*_*x*_ > 1 + *θ*_*y*_, regardless of viral payoff parameters. In this case, all exterior fixed points exhibit saddle stability, which leads us to investigate the existence of heteroclinic cycles.

**Table 8:** Eigenvalues for exterior fixed points of nutrient-explicit model.

| $x$ | $y$ | $n$ | $\lambda_1$ | $\lambda_2$ | $\lambda_3$ |
| --- | --- | --- | --- | --- | --- |
| 0 | 0 | 0 | $b - d$ | $\gamma - \delta$ | -1 |
| 0 | 0 | 1 | $b' - d'$ | $\gamma' - \delta'$ | 1 |
| 0 | 1 | 1 | $a' - c'$ | $-(\gamma' - \delta')$ | $1 + \theta_y$ |
| 0 | 1 | 0 | $a - c$ | $-(\gamma - \delta)$ | $-(1 + \theta_y)$ |
| 1 | 0 | 0 | $-(b - d)$ | $\alpha - \beta$ | $\theta_x - 1$ |
| 1 | 0 | 1 | $-(b' - d')$ | $\alpha' - \beta'$ | $-(\theta_x - 1)$ |
| 1 | 1 | 1 | $-(a' - c')$ | $-(\alpha' - \beta')$ | $-(\theta_x - (1 + \theta_y))$ |
| 1 | 1 | 0 | $a - c$ | $-(\alpha - \beta)$ | $\theta_x - (1 + \theta_y)$ |

#### 5.3.2 Solving Characteristic Matrix

Hofbauer’s [18] strategy for characterizing heteroclinic cycles in replicator dynamics involved a transformation of the equations into a finite number of half-spaces and evaluating the eigenvalues at their intersections. Our model can be redefined in terms of half-spaces in ℝ^3^:

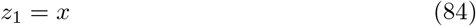

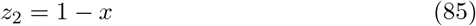

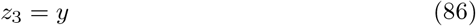

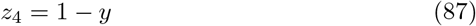

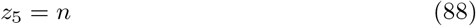

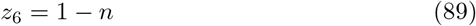

Implementing this rearrangement on the system of differential equations yields:

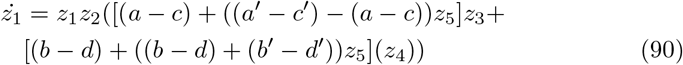

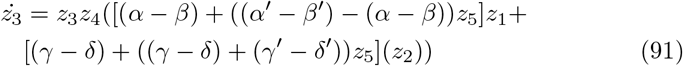

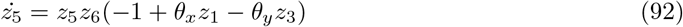

The stability of each plane can be evaluated for fixed points which exist on those planes per the demonstration in [18]. For example, determining whether the equilibrium *x, y, n* = 0 is stable on the plane *x* = 0:

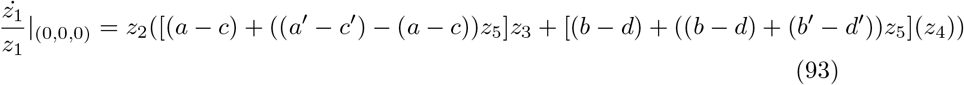

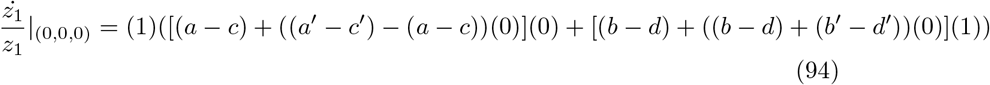

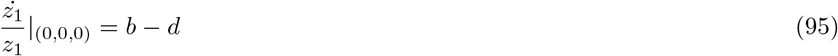

If *b* − *d* is positive, *x, y, n* = 0 is repelling from the plane *x* = 0. Applying this strategy to all fixed points for all directions, we get the following matrix:

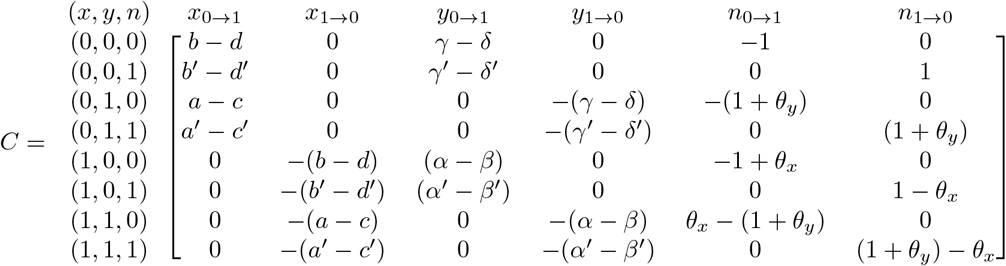

We can read the stability of each exterior fixed point in the directions of its adjacent fixed points off of this matrix. If there are any such set of vertices that form a closed loop with each other, this is a heteroclinic cycle. In the case such that are are more than one closed loops, then multiple possible heteroclinic cycles emerge, and this is called a heteroclinic network. If each row had exactly one positive and one negative value, then the system would have a simple heteroclinic cycle [18]. Using the dynamics demonstrated in Figure 3, we will work through an example characterizing this heteroclinic cycle.

We use the payoffs:

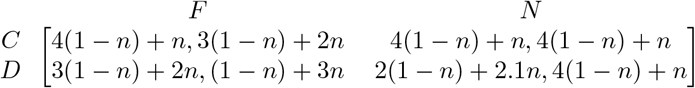

and the environmental restoration parameters *θ*_*x*_ = 4, *θ*_*y*_ = 1. Plugging these in to *C*, the resultant characteristic matrix is:

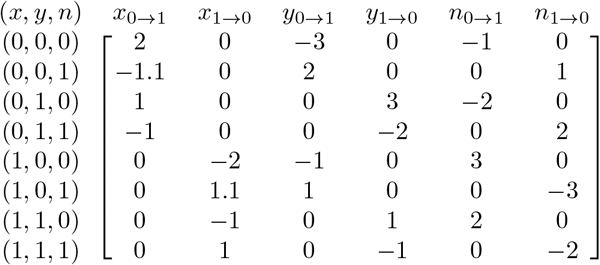

Each row of this matrix has at least one positive value and one negative value, and four of the rows have more than one positive value. This means that a heteroclinic network exists. To identify a cycle, start at any given point and follow the directions indicated by the positive eigenvalues. This network is comprised of the following cycles:

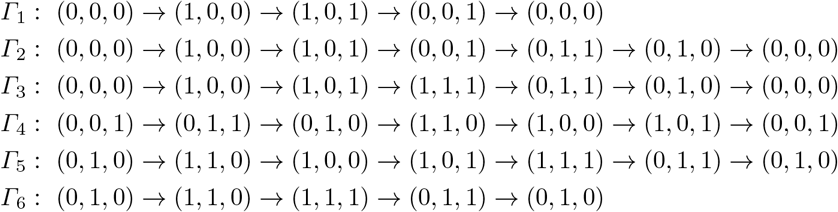

Now we investigate the relative stability properties of each of these cycles. Because all of these cycles contain at least one positive transverse eigenvalue (there are 2 positive values for some row in *C* for at least one vertex in each cycle), none of these cycles will be asymptotically stable. Brannath in [6] describes heteroclinic cycles as essentially asymptotically stable if, aside from some cusp shaped region of initial conditions with small Lebesgue measure, trajectories are attracted to the cycle. For this heteroclinic network, we use the two necessary and sufficient conditions for essential asymptotic stability from [23] 4.11:

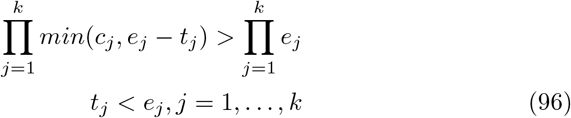

Where *e*_*j*_ is the eigenvalue in the direction of the cycle leaving the fixed point *j* (expanding eigenvalue), *c*_*j*_ is the absolute value of the eigenvalue in the direction of the cycle arriving at fixed point *j* (contracting eigenvalue), and *t*_*j*_ is the eigenvalue in the direction away from the cycle (transverse eigenvalue). *t*_*j*_ can take on positive or negative values. We can calculate these stability criteria for any cycle in the heteroclinic network. Take *Γ*_1_ as an example. These eigenvalues are:

| Point | $-c$ | $e$ | $t$ |
| --- | --- | --- | --- |
| (0,0,0) | -1 | 2 | -3 |
| (1,0,0) | -2 | 3 | -1 |
| (1,0,1) | -3 | 1.1 | 1 |
| (0,0,1) | -1.1 | 1 | 2 |

We find *t*_(0,0,1)_ > *e*_(0,0,1)_, and conclude *Γ*_1_ is unstable. Because all heteroclinic cycles in the network are connected by at least one fixed point, there will only be one cycle for which there is no *t*_*j*_ > *e*_*j*_. For these payoffs, that cycle is *Γ*_2_. We then conclude that any trajectory which is attracted to this heteroclinic network will be attracted to *Γ*_2_. Also notice for *Γ*_1_ that *c* = *e*. This will be the case for all heteroclinic networks in our model because of the symmetrical nature of the eigenvalues. This means that no heteroclinic cycle in our model will be essentially asymptotically stable. Simulations find that without an unstable interior fixed point to drive dynamics towards the exterior of the state space, they dynamics tend to the neutrally stable periodic orbits on the *y* = 0, 1 planes (see Figure 8). This result demonstrates that the neutral 2D orbit outcome (in which one viral type reaches fixation) can occur for parameter regimes outside of a dominating strategy for a particular viral phenotype.

